# Goat genome assemblies, nine breeds for pangenome contribution

**DOI:** 10.64898/2026.10.08.757565

**Authors:** Birbes Clément, Eché Camille, Vernette Caroline, Marcuzzo Camille, Sarry Julien, Suin Amandine, Kuchly Claire, Rogier Odile, Gaspin Christine, Milan Denis, Donnadieu Cécile, Iampietro Carole, Klopp Christophe, Tosser-Klopp Gwenola

## Abstract

The goat research community benefits from one of the most comprehensive livestock genome assemblies, ARS1. However, the rich diversity of goat breeds suggests that a single individual genome is not enough to represent the species full genomic content. Third-generation sequencing technologies have recently made it possible to generate very high quality assemblies that should enable to elucidate unique and/or complex phenotype determinisms depending on genetic structural variations.

In the SeqOccin project, nine goat breeds were selected to represent French goat genetic diversity: Alpine, Boer, Créole, Fossés, Poitevine, Provençale, Pyrénéenne, Rove and Saanen. High-molecular-weight DNA was extracted from blood samples and subjected to long read whole-genome sequencing. There reads were used to generate *de novo* assemblies and were aligned onto the public reference to call structural variants. PacBio HiFi and parental Illumina reads were used to assemble phased haplotypes of the Alpine breed. For the other breeds, PacBio raw (CLR) or ONT reads were generated. Our work demonstrates the feasibility of assembling genomes with a reasonable amount of data and appropriate software packages, delivering one high-quality assembly and 8 standard goat assemblies that pave the way for pangenome construction and structural variant functional studies.

## Introduction

Goats (Capra hircus) were first domesticated around 10,500 years ago in the Fertile Crescent (Zeder, 2008). Following their dispersal from their center of domestication, goats underwent intense adaptation and now occupy diverse agroecological zones around the world. As a result of selection, different goat breeds and lines are specialised for milk, meat and fiber production (Skapetas & Bampidis), as well as in the production of fuel or fertilizer from manure. Thus goats play an important role in the global livestock sector. According to FAOSTAT (www.fao.org/faostat/en/#home), the global goat population has increased by 67% since 1994, surpassing 1.25 billion heads in 2023.

In France, 1,275,000 goats are raised (www.fao.org/faostat/en/#home), mainly for milk and cheese production. Today, France officially recognises fourteen goat breeds, 11 of which are located in metropolitan France and three of which are found in French overseas regions. Ninety percent of the French goats belong to the Alpine or the Saanen breeds. These two industrial dairy breeds originate from eastern France and Switzerland and are managed within efficient breeding programs (Danchin-Burge et al., 2012). The remaining stock is mostly made up of local dairy breeds of moderate size (about 1,000 individuals): the Corse, Poitevine and Pyrénées breeds. Finally, two local breeds with fewer than 1,000 individuals are managed within conservation programs: the Provençale and the Fossés breeds. In addition to these dairy breeds, a small population of Angora individuals exists that is reared for Mohair production. This breed originates from Turkey and is managed in an organized breeding program (Danchin-Burge et al., 2012).

In 2010, the first goat reference genome was sequenced and assembled (Dong et al., 2013), and the International Goat Genome Consortium (IGGC) was created to support the development of genomic tools for studying the genetic variation of domestic goats (www.goatgenome.org) The next major step was the development of the GoatSNP50 BeadChip in 2014 (Tosser-Klopp et al., 2014), a 50 K single nucleotide polymorphism (SNP) panel. The GoatSNP50 chip enabled the discovery of quantitative trait locus (QTL) through genome-wide association studies (Martin et al., 2016, 2017; Mucha et al., 2018) and helped to initiate genomic selection in goats (Carillier et al., 2013; Carillier-Jacquin et al., 2016). It also allowed both global and more specific characterisation of goat genetic diversity. The AdaptMap dataset collected SNP chip data from 4,653 animals across 148 populations and 35 countries (Stella et al., 2018). For French breeds, the AdaptMap data were extended to other French breeds in Oget et al. study (Oget et al., 2019). The goat assembly was updated in 2017 with ARS1 (GCA_001704415.1), which is the current Ensembl reference genome (Bickhart et al., 2017) and SNP chips have been either updated or created (www.goatgenome.org).

However, data generated from SNP chips are known to be distorted by ascertainment bias (Albrechtsen et al., 2010), a limitation that can be overcome by carefully filtering whole-genome sequencing data of sufficient depth (Benjelloun et al., 2019). This is one of the reasons why large-scale short-read sequencing projects have been constructed, like the VarGoats project, that has generated new data and gathered public data to detect variants from 1,159 genomes (Denoyelle et al., 2021) as a first step and 1,372 genomes in a second step (www.goatgenome.org/vargoats).

One of the limitations of such studies is the dependence on the reference assembly. For example, large-size insertions can hardly cannot be detected through the alignment of short-reads on a reference assembly.

Recently, long-read sequencing technologies have enabled the construction of *de novo* assemblies with a lower cost. In one of its work packages, the SeqOccin project aimed at generating assemblies in a set of species, among which *Capra hircus*. Several French breeds were chosen to this extent. Nine animals were selected in 9 breeds (Alpine, Boer, Créole, Fossés, Poitevine, Provençale, Pyrénéenne, Rove and Saanen) to represent the goat genetic diversity in France. Alpine and Saanen were chosen as they represent 90 % of the French goats animals. Boer and Creole breeds were chosen for their “overseas origin” while the Rove breed was chosen for specific traits (horns). The other breeds were selected as (Oget et al., 2019) showed that they differ in terms of genetic background.

In this paper, we deliver one high-quality assembly and eight standard assemblies in nine French goat breeds, and we expect them to serve pangenome construction projects and help elucidate complex phenotypes through the identification of structural variants.

## Material and methods

### Genome information

The domestic goat genome follows the *Bovidae* family chromosomal organisation with 29 autosomes and 2 sex chromosomes (2n = 60). As in other *Bovidae*, autosomes are acrocentric with terminal centromeres. Genome size estimations for goat range from 2.66 Gb to 3.04 Gb, depending on the method (flow cytometry, Feulgen staining or sequencing-based analyses). Based on k-mer analysis (Marçais & Kingsford, 2011; Ranallo-Benavidez et al., 2020) with PacBio HiFi data, the goat genome is estimated at ~2.92 Gb with heterozygosity levels around 0.25– 0.30% and ~36% repeated content.

The current NCBI ARS1.2 (GCA_001704415.2) reference assembly metrics are: 2.92 Gb total size, 23.6 Mb contig N50, 87.3 Mb scaffold N50, and 97.5% BUSCO score. We used ARS1.2 as the reference to evaluate the quality of our assemblies.

### Sample collection & technical validation

Blood samples were collected by the GenPhySE unit, as allowed by the government under the agreement number: 31113050. The Qiagen Tips 100 was used to extract high molecular weight DNA from these goat fresh blood samples. The extracted DNA was stored at ™20°C in TE buffer (10 mM Tris–Cl, pH 7.5; 1 mM EDTA)

For Alpine parental short-read sequencing, an in-house protocol was used. Genomic DNA concentrations were measured using the Qubit fluorimetry system (Life Technologies) with the High Sensitivity (HS) kit for detection of double-stranded DNA (Thermo Fisher, Part #Q32854). Fragment size distributions were assessed using the Femto pulse DNA 165kb Kit (Agilent). Purity was measured using a Nanodrop system (Thermofisher). All the samples were purified with AMPureXP beads (Beckman Coulter, A63882) to obtain correct ratios *i*.*e*. 260/280: 1.8–2 and 260/230:2-2.2.

### Sequencing

All sequencing processes were performed at the GeT-PlaGe core facility at INRAE Toulouse (GeT-PlaGe, 2022).

At each step, DNA was quantified using the Qubit HS kit. DNA purity was tested using a Nanodrop and size distribution and degradation assessed using the Femto pulse DNA 165kb Kit or Fragment Analyser kit. Purification steps were performed using AMPure PB beads (PacBio 100-265-900).

#### PacBio Sequel II

Library preparation was performed according to the manufacturer’s instructions “Procedure & Checklist Preparing HiFi SMRTbell Libraries using SMRTbell Express Template Prep Kit 2.0” (100-938-900).

For CLR library (simple assembly): 10µg of DNA were purified then sheared at 60kb using the Megaruptor3 system (Diagenode). Using SMRTbell Express Template prep kit 2.0, a Single strand overhangs removal, a DNA and END damage repair step were performed on 10µg of sample. Then, blunt hairpin adapters were ligated to the library. A size selection step using a 30kb cutoff was performed on the BluePippin Size Selection system with “0.75% DF Marker U1 high pass 30–40kb VS3” protocol. A 70kb library was recovered. Using Binding kit 2.0 kit (101-789-500) and sequencing kit 2.0 (101-820-200) with 1 h of annealing and 4h of binding, the primer V4 annealed and polymerase 2.0 bounded library was sequenced by diffusion loading onto 1or 2 SMRTcells 8M (101-389-001) on Sequel II instrument at 50 to 70 pM with a 15-hour movie. A coverage of minimum 30X was obtained per sample.

For HiFi library (Alpine assembly), 20µg of DNA were purified then sheared at 15kb using the Megaruptor3 system (Diagenode, N°E07010003). Using SMRTbell Express Template prep kit 2.0, a Single strand overhangs removal, a DNA and END damage repair step were performed on 10 µg of sample. Then blunt hairpin adapters were ligated to the library. The library was treated with an exonuclease cocktail to digest unligated DNA fragments. A size selection step using a 10kb cutoff was performed on the BluePippin Size Selection system to obtain a 20kb library. Using Binding kit 2.0 kit and sequencing kit 2.0 with 1h for annealing and 1h for binding, the primer V2 (101-847-900) annealed and polymerase 2.0 bounded library was sequenced by diffusion loading onto 4 SMRTcells on Sequel II instrument at 50 to 70 pM with a 2 hours pre-extension and a 30-hour movie. A coverage of minimum 40X was obtained.

#### Illumina paired-end WGS 2 × 150 bp

To generate the high-quality Alpine assembly, the parents were sequenced as follows. DNA-seq libraries were prepared according to the Illumina protocol TruSeq Nano DNA High Throughput Library Prep Kit. DNA was fragmented by sonication on Pixul, size selection was performed using Sample Purification Beads of the library kit (ratio 1/1 beads water) and adaptators IDT for Illumina – TruSeq DNA UD Indexes were ligated before sequencing. Library quality was assessed using a Fragment Analyzer (Agilent) with High Sensitivity NGS Kit (DNF-474-0500). Sizes of 600 bp were obtained. Libraries were quantified by qPCR using the KAPA Library Quantification Kit (Roche, 07960140001). DNA was sequenced on 1/2 S4 NovaSeq 6000 lane using a paired-end read length of 2 × 150 bp with the Illumina S4 Reagent kit (300 cycles). This sequencing produced a 70X coverage per sample.

#### Hi-C

Blood sample was processed using the Arima Hi-C Kit (ref. 510008) according to the manufacturer’s protocols. The library was prepared using the TruSeq DNA PCR-Free Kit and quantified by qPCR on QuantStudio 6 device (Applied Biosystems, ThermoFisher Scientific) using the Kapa Library Quantification Kit (Roche, KK4824). Sequencing was performed in a 2 × 150 paired-end mode on a NovaSeq6000 (Illumina).

#### ONT

At each step, the DNA was quantified using the Qubit HS kit. DNA purity was tested using the nanodrop and size distribution and degradation were assessed using the Fragment analyzer (AATI) High Sensitivity DNA Fragment Analysis Kit. Purification steps were performed using AMPure XP beads (Beckman Coulter). Library preparation was performed according to the manufacturer’s instructions “1D gDNA selecting for long reads (SQK-LSK114)”. For 1 Flowcell, 5µg of DNA were purified then sheared at 20 kb, using the Megaruptor system (Diagenode). A 10kb size selection step using the BluePippin Size Selection system was performed. A one-step DNA damage repair + END-repair + dA tail (NEB) of double stranded DNA fragments was performed on 1µg of sample. Then, adapters were ligated to the library. The library was loaded onto FLO-PRO004 (R10.4.1) flowcells and sequenced using PromethION instruments at 6/10 fmol within 72h. Nuclease flush steps were done when necessary and possible, *i*.*e*. when only 30% of the pores were still in sequencing. We produced a 20 to 50X coverage.

### Genome assembly

As presented in the previous section, several types of reads were produced during the project. Three types of long reads were generated: PacBio CLR for Boer, Créole, Fossés, Poitevine, Provençale, Pyrénéenne, Rove and Saanen breeds, PacBio HiFi for the Alpine breed and Oxford Nanopore for the Créole breed. For the Alpine animal only, Hi-C reads were used for scaffolding and Illumina short reads were used for phasing chromosomes pairs. The long read types were differing in error rate and therefore could not be assembled using a unique piece of software. Because of initial low throughput for Créole, ONT reads were produced two years later and a third up-to-date assembly strategy was used.

The first « High quality » strategy, used for the Alpine breed only, was based on HiFi reads assembled with hifiasm version 0.16.1 (Cheng et al., 2021) and parental kmer dictionaries (−1 and −2 parameters) to separate haplotypes on the chromosome level. The kmer dictionaries were produced with yak (Li, 2026) count version 0.1 and a kmer size of 37. The haplotype separation was checked with merqury version 1.3 (Rhie et al., 2020) using default parameters. Before running merqury, parental reads were used to produce kmer dictionaries with meryl version 1.3 (Miller et al., 2008) with default parameters. The second “Medium quality” assembly strategy was based on PacBio CLR reads which were assembled for six breeds with flye version 2.9 (Kolmogorov et al., 2020) with the --pacbio-raw and --genome-size 3.0g parameters. Because of the low read quality, this strategy includes a contig polishing round using the same long reads which was performed with ArrowGrid-release 489ebf8 from SMRTLink version 10.2 (Chin et al., 2013). Briefly, the reads were realigned onto the contigs and the software package checked the sequencer signal of all reads for all positions to decide which nucleotide to put into the consensus.

Because the Créole DNA sample did not sequence well with PacBio CLR, a new DNA preparation was sequenced with Oxford Nanopore Technologies. In this third strategy CLR plus ONT reads were assembled with pecat (Nie et al., 2024; lemene, 2026) unzip version 0.0.3 using default parameters which first corrected the input reads by aligning them all versus all. These corrected reads were then assembled and separated into two haplotypes which were polished independently. This strategy produced two haplotypes per breed.

### Genome scaffolding

Two scaffolding strategies were used. The first one was used for the Alpine breed, based on Hi-C reads in which both haplotypes were scaffolded independently. First, the Hi-C reads were aligned to each haplotyped contig set with juicer version 1.5.6 (Durand, Shamim, et al., 2016) with default parameters. The resulting Hi-C link file was processed with 3d-dna release 529ccf4 with parameter –s 0 in order to stop scaffolding at the first round. The resulting scaffolding was manually curated using juicebox version 1.11.8 (Durand, Robinson, et al., 2016) to produce chromosomes. Chromosomal assemblies were checked with merqury.

All the other breed assemblies were scaffolded on the ARS 1.2 reference genome from the NCBI website (https://ftp.ncbi.nlm.nih.gov/genomes/all/GCF/001/704/415/GCF_001704415.2_ARS1.2/GCF_001704415.2_ARS1.2_genomic.fna.gz) with ragtag version 2.0.1 using minimap2 version 2.24 (Li, 2018). Briefly, after aligning the contigs to the reference chromosomes, the alignment uniqueness and quality enabled to decide contig location and orientation and contigs were then merged and separated in chromosomes with a spacing sequence of 1000 Ns between consecutive contigs. Non-matching or non-uniquely matching contigs were kept isolated in the resulting fasta file. Pyréennne, Poitevine and Saneen assemblies showed two large missassemblies and one large mis-scaffolding - one in each assembly - which were manually corrected.

### Assembly validation

Several metrics were used to validate the scaffolded assemblies. For all of them assemblathon_stats.pl script was run to extract contig and scaffold statistics including total length, N50, L50 and GC content. Busco scores (Manni et al., 2021) were also produced in all cases, using version 5.1.2 with mammalia_odb10 database. For high quality and corrected reads, kmer QV and completeness were calculated using merqury (Rhie et al., 2020) version 1.3 with default parameters and a blob graph was generated for the Alpine breed, using parental reads kmer content.

## Results

The data presented hereunder has been used to produce high-quality phased genome assemblies for the Alpine breed, one haplotyped assembly for creole, and 7 assemblies for the Fossés, Poitevine, Provençale, Pyrénéenne, Rove and Saanen breeds.

The metrics below highlight the high quality of these data.

**Table 1:** Goat data volume. Data volume for each breed. The first column corresponds to the goat breed, the second to the number of reads, the third to the total read size, the fourth to the average read length, the fifth to the sequencing coverage, and the last one to the EBI/EMBL identifiers.

| Breed | Read numbers | Total size | Average length of reads | Coverage (X) | ID EBI/EMBL |
| --- | --- | --- | --- | --- | --- |
| Alpine | 5 413 668 | 94 408 246 598 | 17 439 | 32 | ERR14939006, ERR14939010, ERR14939011,ERR14939012, ERR15074730 |
| Boer | 4 215 057 | 165 294 842 127 | 39 215 | 57 | ERR14939007 |
| Créole | 21 228 058 | 256 847 142 116 | 12 099 | 89 | ERR14939037, ERR14939039, ERR15094489 |
| Fossés | 6 942 204 | 141 491 052 628 | 20 381 | 49 | ERR14939008 |
| Poitevine | 6 942 204 | 141 491 052 628 | 20 381 | 48 | ERR14939002 |
| Provençale | 10 159 681 | 216 817 630 122 | 21 340 | 75 | ERR14939003, ERR14939004, ERR14939005 |
| Pyrénéenne | 15 275 217 | 327 691 808 292 | 21 453 | 113 | ERR14938999, ERR14939000, ERR14939001 |
| Rove | 6 124 116 | 225 982 780 710 | 36 900 | 78 | ERR14939009 |
| Saanen | 3 553 782 | 139 667 911 503 | 39 301 | 48 | ERR14939013 |

**Table 2:**
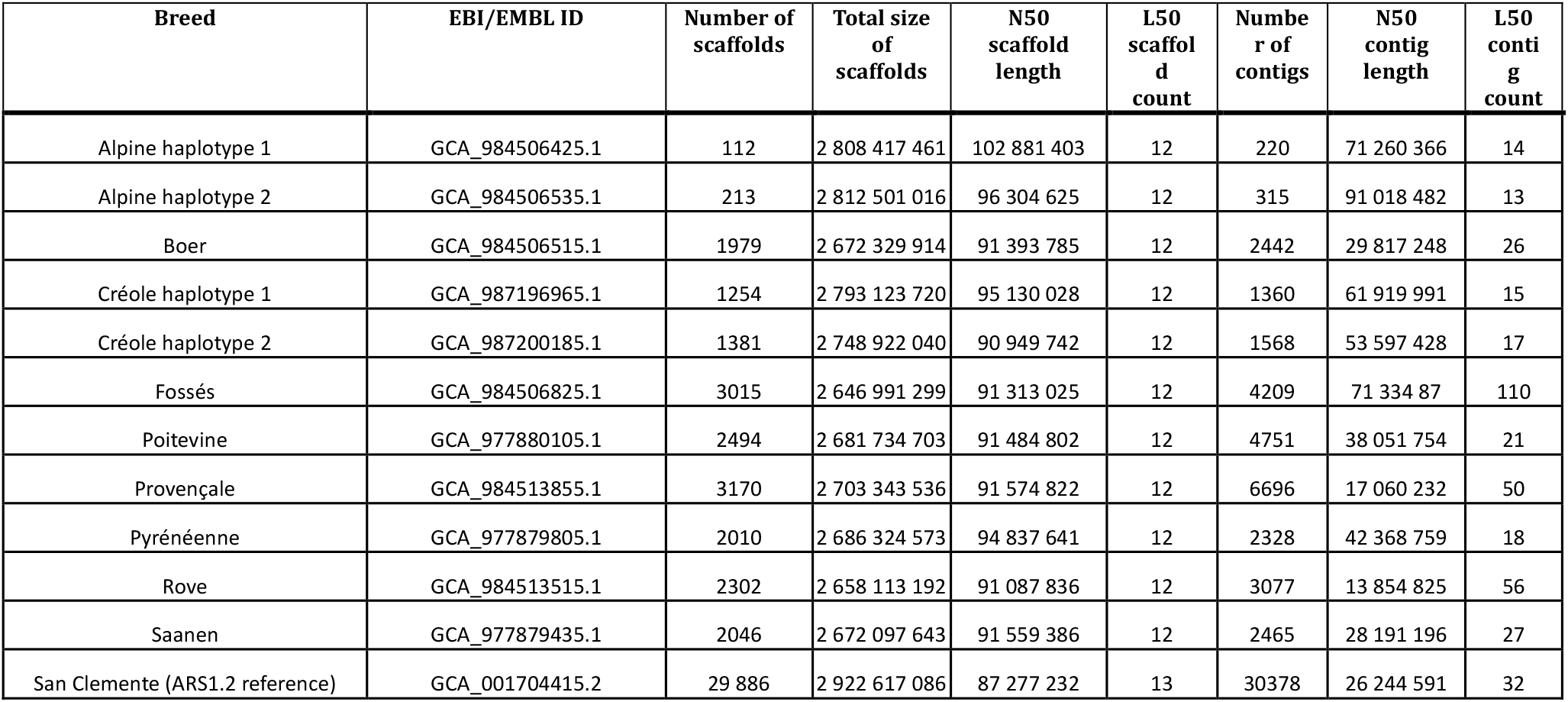
Goat Assembly statistics. Evaluation per breed of assembly quality and comparison with the reference (GCF_001704415.2_ARS1.2_genomic.fna.gz). Column 1 gives the breed (and haplotype if present), column2 the corresponding EBI/EMBL ID, column 3, the number of scaffolds obtained using the assembler, column 4, the total assembly length, column 5 and 6, the scaffold N50 and L50. Columns 7,8,9 correspond to the metrics generated on contigs.

**Table 3:**
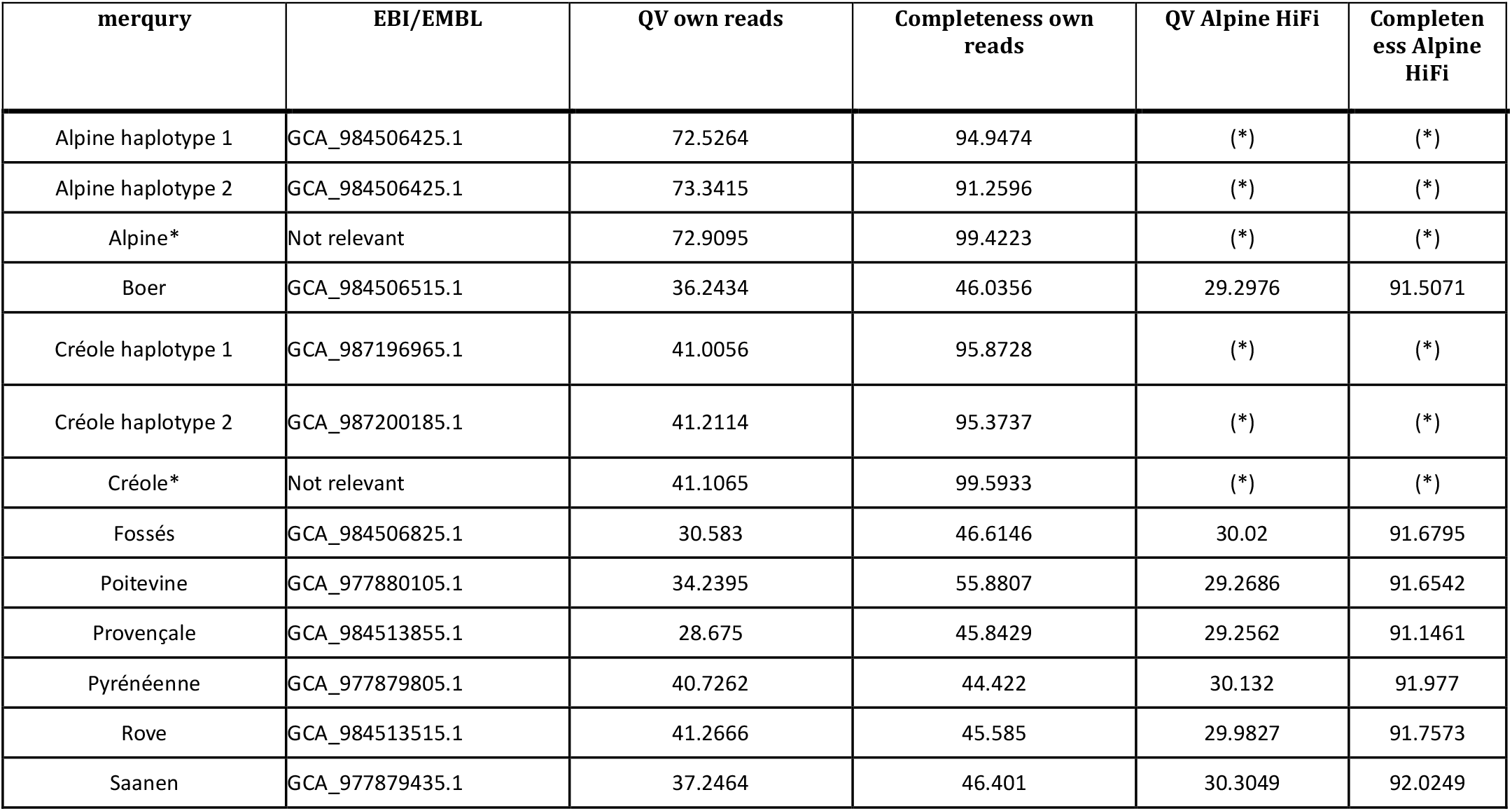
Merqury. (*)Not calculated for HiFi and ONT reads For each assembly the table presents merqury QV and completeness results. Column 1 gives the breed (and haplotype if present), column2 gives the assembly ID, column 3 and 4 present QV and completeness calculated using the reads with which the assembly was performed. Because CLR reads are of low quality (close to 12% errors) these metrics are poor. Therefore column 5 and 6 present the same metrics, but calculated with the Alpine HiFi reads. (*) Both previous haplotypes were used.

**Figure 1.**
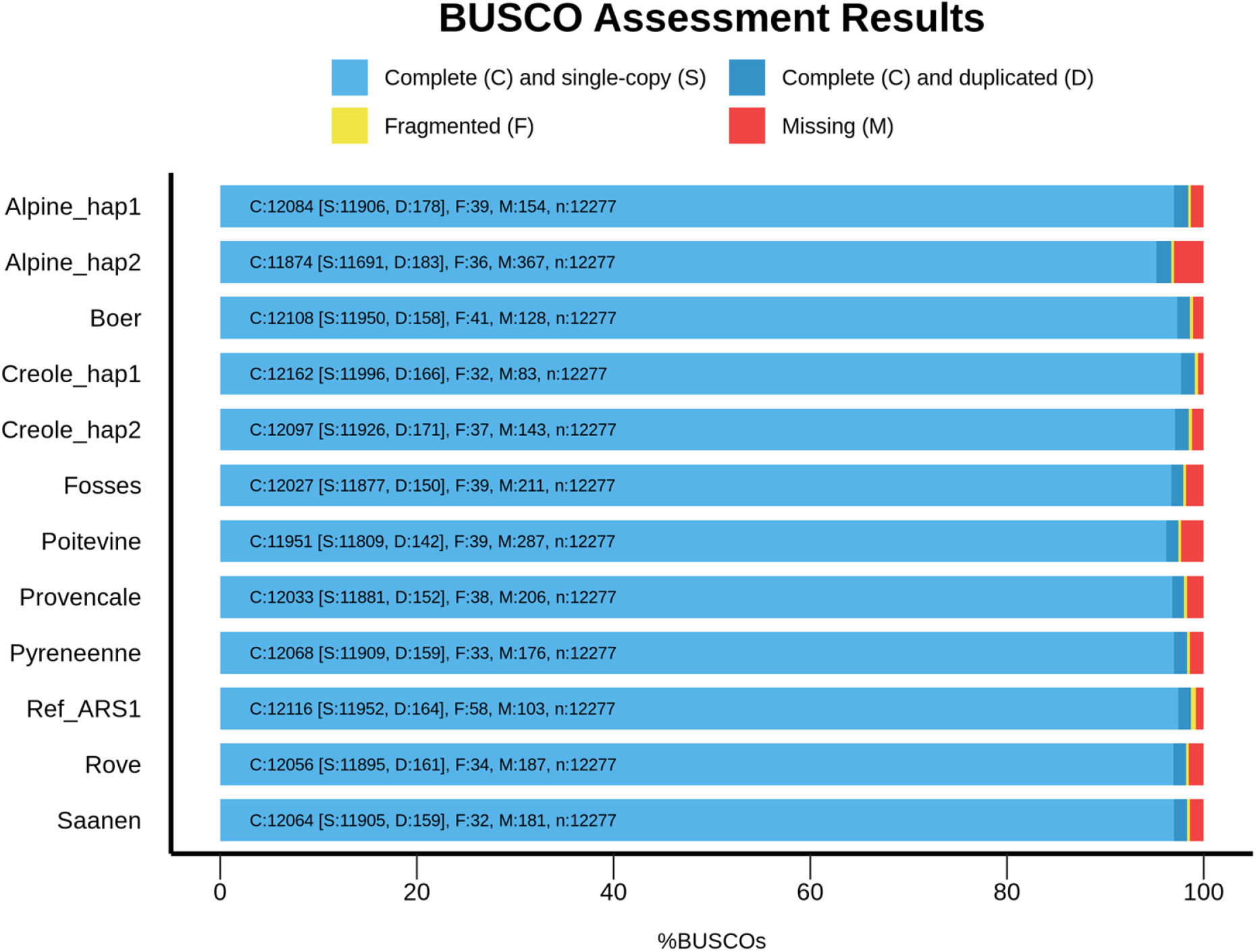
BUSCO. BUSCO plot of the goat assemblies. Summary plot produced with BUSCO V5.1 and mammalia_odb10 dataset. The breed is indicated in line and the haplotype is added if relevant (_hap1 for haplotype 1 and _hap2 for haplotype 2). Ref_ARS1 is the ARS1.2 reference assembly.

## Acknowledgements

We thank all the people, who selected the breeds and collected the blood samples. The following list is probably not exhaustive:

- Isabelle Palhière and other researchers, engineers and technicians from the Unité Mixte de Technologie STAR (https://idele.fr/umt-star/), Coralie Danchin-Burge (IDELE).
- technicians of INRAE experimental units for the following breeds: Alpine (P3R – Pôle de phénotypage des petits ruminants (INRAE Unité Expérimentale Pôle de Phénotypage des Petits Ruminants Bourges (P3R Bourges) Domaine de la Sapinière 18390 Osmoy – France, https://doi.org/10.15454/1.5483259352597417E12) and Créole (*PTEA*, INRAE, 2026, Plateforme Tropicale Élevage pour l’Agroécologie https://doi.org/10.17180/50n1-kn86)
- Apolline Bailly-Salins, Capgenes, https://www.capgenes.com/ for Boer, Provençale and Saanen breeds
- the breeders and the relevant breeding organisations for Fossés, Poitevine, Provençale, Pyrénéenne and Rove breeds.

We thank “La Région Occitanie” and the European Union for funding the project as part of the Occitanie Region’s “Regional Research and Innovation Platforms” call for projects under the FEDER-FSE MIDI-PYRENEES ET GARONNE 2014-2020 Operational Program.

GeT core facility (https://doi.org/10.15454/1.5572370921303193E12) is supported by France Génomique National infrastructure, funded as part of “Investissement d’avenir” program managed by the French Agence Nationale pour la Recherche (contract ANR-10-INBS-09). We are grateful to the genotoul bioinformatics platform Toulouse Occitanie (Bioinfo Genotoul, https://doi.org/10.15454/1.5572369328961167E12) for providing computing and storage resources.

## Data, scripts, code, and supplementary information availability

Data are available online:

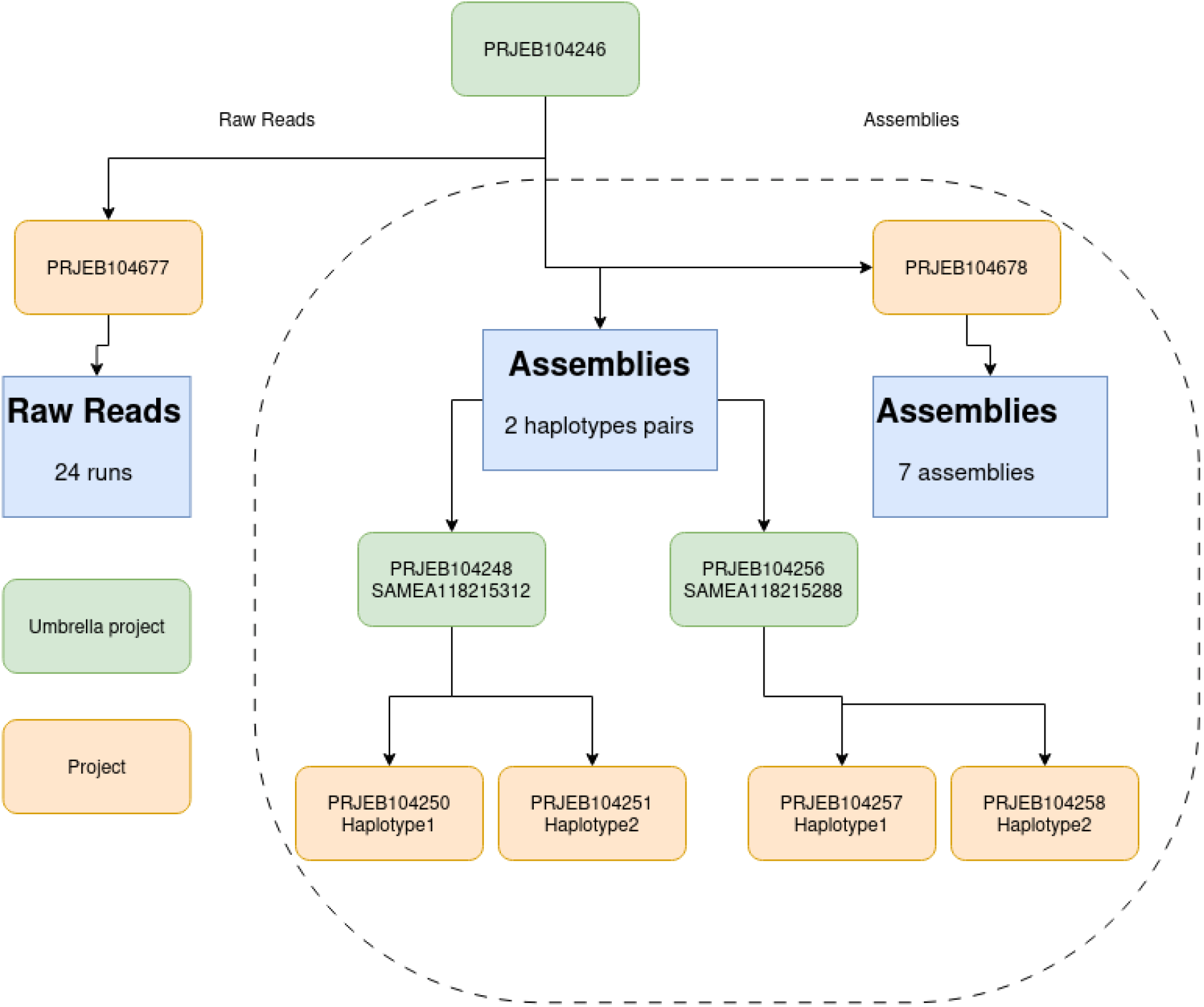

SWHID for the software packages used in this project. These IDs enable to download the software packages from https://archive.softwareheritage.org/

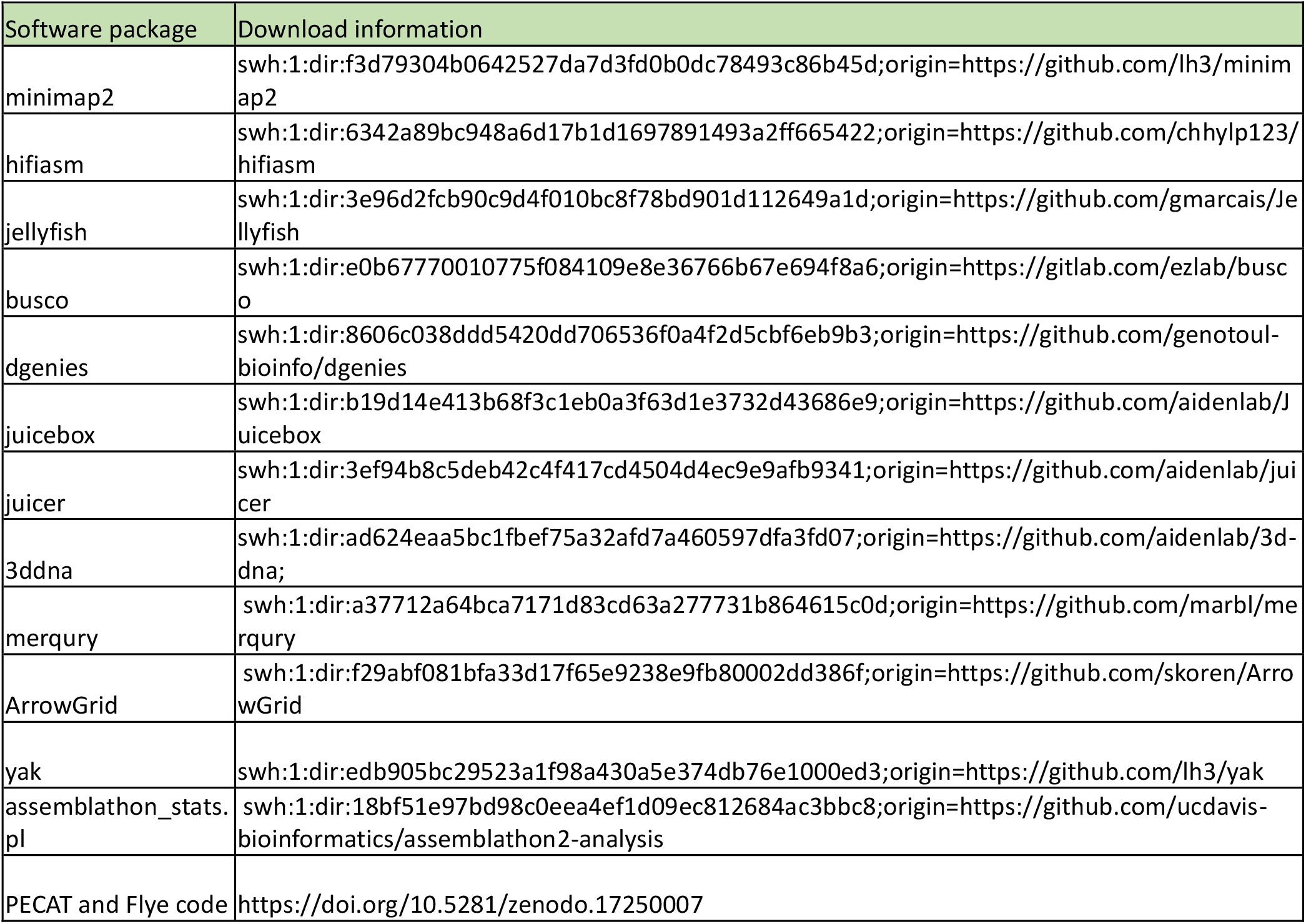

